# Sexually dimorphic neural encoding of threat discrimination in nucleus accumbens afferents drives suppression of reward behavior

**DOI:** 10.1101/2022.10.04.510865

**Authors:** J Muir, YC Tse, E Iyer, J Sorensen, R Eid, V Cvetkovska, K Wassef, S Gostlin, NJ Spencer, RC Bagot

**Author notes:** Authors contributed equally to this work (YC Tse, E Iyer).

## Abstract

Learning to predict threat is essential but equally important yet often overlooked is learning about the absence of threat. Interrogating neural activity in two nucleus accumbens afferents in male and female mice during aversive and neutral cues reveals sex-specific encoding of cue discrimination and cue-mediated suppression of reward behavior. Sexual dimorphisms in neural bases of threat discrimination may reflect sex differences in behavioral strategy relevant to differential psychiatric risk.

## Main Text

The importance of learning to recognize and respond to impending threat is widely appreciated but the related process of recognizing when threat is absent is commonly overlooked. Threat inhibits appetitive behavior (e.g., food seeking) avoiding risk but by learning when threat is not imminent, an animal can pursue essential goals in periods of relative safety. Neural mechanisms discriminating aversive and neutral events underpin adaptive behavior and are disrupted in psychopathology ^1-3^. The nucleus accumbens (NAc) integrates diverse inputs ^4-6^, balancing threat and reward to orchestrate motivated behavior ^6-9^. Glutamatergic projections from the ventral hippocampus (vHip) and medial prefrontal cortex (mPFC) to NAc are implicated in reward processing ^4^ and adaptation to chronic stress ^10,11^. How these pathways integrate aversive information to modulate behavior is not fully understood, and, in females, largely unstudied, despite known sex-differences in stress-related psychopathologies ^12,13^. Here, we examined how mPFC and vHip projections to NAc medial shell (mPFC-NAc, vHip-NAc), a major target for both vHIP and PFC afferents, encode aversive experiences to guide behavioral responding to threat.

To probe pathway-specific neural encoding of aversive cues in mPFC-NAc and vHip-NAc, we injected retrograding AAV-GCaMP7f in NAc and implanted optic fibers in vHip and mPFC to record Ca^2+^-associated fluorescence while male and female mice encountered cue-shock (CS+) and cue-no outcome (CS-) pairings (Figure 1a,b). Males and females learned cue-shock associations by mid-training (Figure 1c,d), increasing freezing during the CS+ and discrimination increased across training (Figure 1e). Marginal sex differences in CS-mediated behavior emerged in late-training: CS-suppressed freezing in females, but not males (Figure 1c,d). Freezing suppression ratios confirmed that, relative to baseline, CS+ increased freezing in males and females but CS-suppressed freezing below baseline only in females (Figure 1f-i), suggesting here females may learn the CS-as a safety signal.

**Figure 1.**
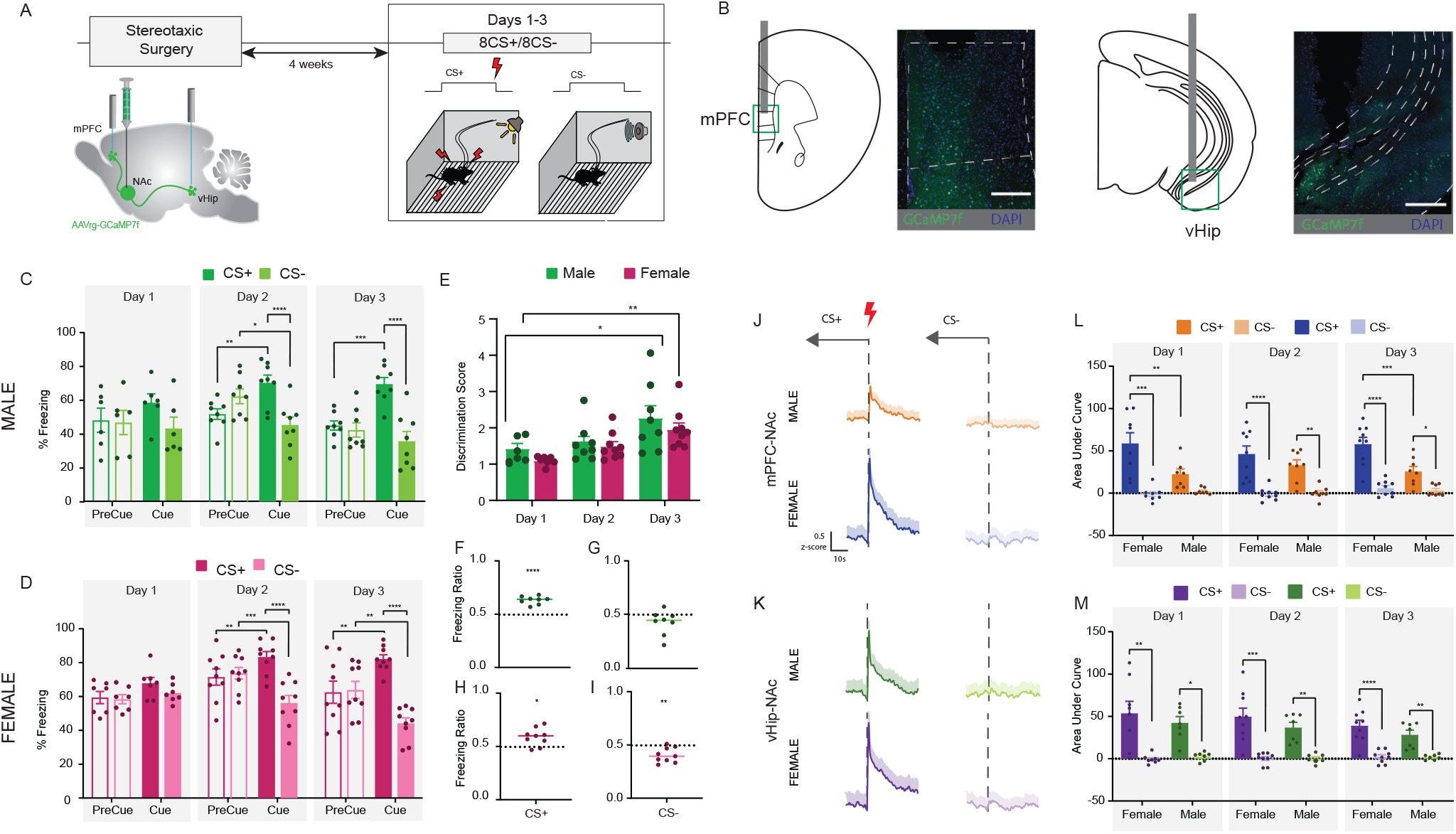
Male and female mice acquire discriminative Pavlovian conditioning. (A) Experimental timeline of surgery and discriminative Pavlovian conditioning. Male and female mice were injected with a retrograding GCaMP7f into the NAc and implanted with fibers above the vHIP and mPFC to image simultaneous projection-specific neural activity in both pathways. Following recovery and time for viral expression, mice were exposed daily to 8 CS+ cues co-terminating with footshock and 8 CS-cues with no outcome for 3 days. (B) Images indicate representative virus and fibre targeting (scale bars represent 250um). (C) In early-training (day 1), males showed a main effect of cue type on freezing (F _(1, 5)_ = 10.32, n=6, p=0.02). By mid-training (day 2), males froze more to the CS+ than to CS-(F _(1, 7)_ = 33.47, n=8, p=0.0007, *post hocs*: p=0.0014) and freezing increased from pre-cue for CS+ (p=0.0074) and decreased for CS- ue (p=0.011). At late-training (day 3) males froze more to the CS+ than CS-(F _(1, 7)_ = 52.08, n=8, p=0.0002, *post hoc*: p<0.0001) and relative to pre-cue, freezing increased during CS+ but was not altered by CS-(p=0.0002). (D) On training day 1, female freezing did not differ by cue type (F _(1, 6)_ = 1.425, n=7, p<0.05). In mid-training, females showed behavioral discrimination, freezing more to CS+ than CS-(F _(1, 8)_ = 85.30, n=9, p<0.0001, *post hocs*: p<0.0001) and, relative to the pre-cue period, increasing freezing to CS+ (p=0.0001) while decreasing freezing to CS-(p=0.0015). At late-training, females continue to discriminate, freezing more to CS+ than CS-(F _(1,8)_ = 54.91, n=9, p<0.0001, *post hocs*: p<0.0001) with CS+ and CS-exerting opposing modulation of freezing relative to pre-cue (p=0.0016). (E) Cue discrimination (% freezing CS+/ % freezing CS-) increased across training in both males and females (F _(2,41)_ = 9.47, n=8,9, p<0.0004, *post hocs*: p=0.0004, p=0.017). Freezing ratio (% freezing during cue/ % freezing during cue + % freezing during pre-cue) on day 3, showed that CS+ significantly increased freezing from pre-cue in (F) males (t_7_=9.85, n=8, p<0.0001) and (H) females (t_8_=3.2, n=9,p=0.012), but CS-significantly suppressed freezing in (I) females (t_8_=4.11, n=9, p=0.003) but not (G) males (t_7_=1.97, n=8, p<0.05). (J,K) Averaged traces (light shading indicates SEM) show that footshock following CS+ increased neural activity in males and females compared to CS-no outcome in (L) mPFC-NAc with females exhibiting larger mPFC-NAc pathway response compared to males (Day 1: F _(1, 14)_ = 8.23, n=8,8, p=0.012, *post hocs*: p_F_<0.0001, p_M_=0.12, p_MvF_=0.0019; Day 2: F_cue (1, 13)_ = 56.74, n=8,8 p<0.0001; F_cuexSex (1, 13)_ = 2.83, n=8,8 p<0.11*post hocs*: p_F_<0.0001, p_M_=0.002, p_MvF_=0.09; Day3: (F _(1, 15)_ = 9.47, n=8,9, p=0.0077, *post hocs*: p_F_<0.0001, p_M_=0.01, p_MvF_=0.0004) and (M) vHIP-NAc projections (Day 1: F _(1, 14)_ = 26.46, n=9,7, p=0.0001, *post hocs*: p_F_=0.0014, p_M_=0.017; Day 2: F _(1, 14)_ = 43.44, n=9,7, p<0.0001, *post hocs*: p_F_=0.0001, p_M_=0.0044; Day3: (F _(1, 14)_ = 50.19, n=9,7, p<0.0001, *post hocs*: p_F_<0.0001, p_M_=0.0029). Graphs show mean +/-SEM.

To understand how threat is encoded, we then examined shock-associated changes in neural activity (Figure 1j,k). Shock increased activity in both pathways (Figure 1l,m), with larger increases in mPFC-Nac in females than males, indicating augmented pathway-specific aversive processing. We next examined CS+ and CS-discrimination encoding (Figure 2a-d). Systematically contrasting CS+ and CS-elicited neural activity using a generalized additive model (Figure 2e) identified two epochs maximally encoding cue identity: 1sec at cue onset and 8sec pre-outcome (Figure 2f-i) and subsequent analyses focus therein. Sex and pathway-specific neural signals emerged across training; activity at cue onset in mPFC-NAc discriminated cue type in females but not males, and in vHip-NAc, in males but not females. In mPFC-NAc in males, a CS-peak emerged in mid-training, with equivalent CS+ and CS-peaks in late-training (Figure 2j,m). In mPFC-NAc in females, CS+ and CS-peaks were similar in early-training with CS+ exceeding CS-peak in late-training (Figure 2k,o). In vHip-NAc, in males, similar CS+ and CS-peaks in early-training resolved to only a CS+ peak in late-training (Figure 2l,p), while in females, CS+ and CS-elicited similar peaks throughout training (Figure 2m,q). We also observed CS+ specific suppression in the pre-outcome period in both pathways and sexes across training (Figure S1).

**Figure 2.**
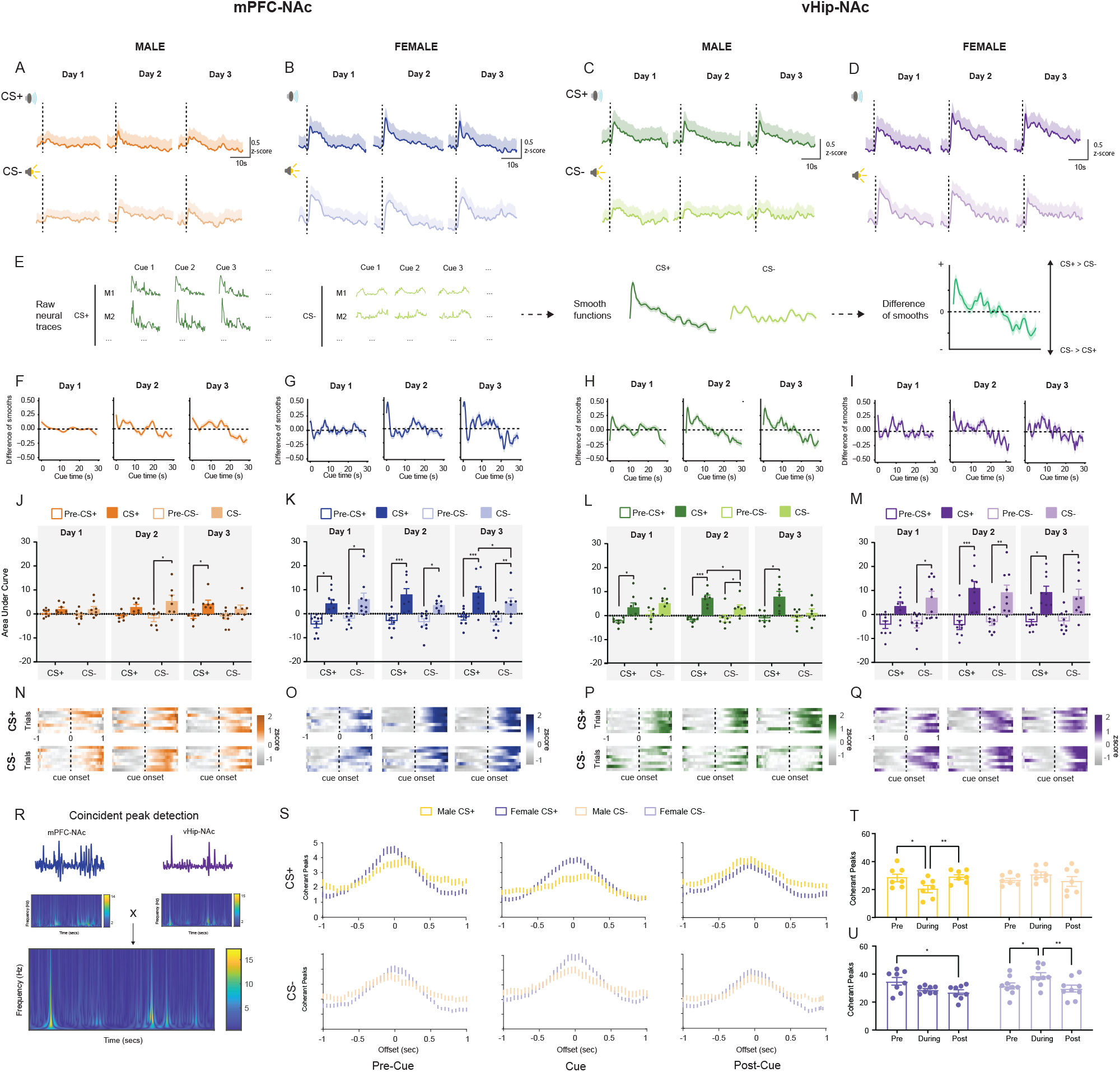
Sex-specific neural encoding of aversive and non-reinforced cues. (A-D) Average time-locked cue-elicited neural activity in mPFC-NAc and vHip-NAc shows peaks for CS+ and CS-(light shading indicates SEM). (E) Generalized Additive Modeling (GAM) uses a sum of smooth functions to model contributions of a fixed variable, cue type, to variation in time series of neural activity recorded during individual cue presentations nested within individual animals, accounting for animal ID as a random variable. To probe differences in CS+ and CS-elicited neural activity across the cue, the difference function of the two smooth functions is calculated, with large non-zero values indicating epochs of maximum difference. (F-I) GAM revealed differences in CS+ and CS-elicited neural activity that emerge across training and identified 2 periods of maximal difference: 1 sec at cue onset and 8 sec preceding cue termination. (J) Further analysis of neural activity at cue onset revealed that in early-training, activity in mPFC-NAc in males was not altered (F _(1,6)_ = 15.44, n=7, p=0.15), in mid-training there was an increase to CS-, but not CS+ (F _(1,6)_ = 20.11,n=7, p=0.004, post hocs: p=0.029) and, during late-training, to CS+ with a trending increase to CS-(F _(1,6)_ = 10.58,n=7, p=0.017, post hocs: p_CS+_=0.013, p_CS-_=0.08) (Figure 3a). (K) In females, PFC-NAc activity increased at cue onset for both cue types in early training (F _(1,8)_ = 24.25, n=9, p=0.0012, post hocs: _CS+_=0.011, p_CS-_ =0.017), and in mid-training, (F _(1,8)_ = 26.92,n=9, p=0.0008, post hocs: p_CS+_<0.0004, p_cs-=_0.012), with increased activity to CS+ relative to CS-in late training (F _(1, 8)_ = 22.5,n=9, p=0.0015, post hocs: p_cue_<0.04, _pre-cue_=0.34) (p=0.036). In contrast, in the vHIP-NAc, (L) in males activity increased at cue onset to both cues in early training (_(1,6)_ = 17.06, n=7, p=0.006, post hocs: p_CS+_=0.02, p_CS-_=0.06) and mid-training (F _(1,6)_ = 58.66,n=7, p=0.0003, post hocs: p_CS+_=0.0004, p_cs-_=0.021) but activity to CS+ was significantly greater than to CS-in mid-training (p=0.025) and by late-training activity was only increased to CS+ (F _(1,6)_ = 13.16,n=7, p=0.01, post hocs: p_CS+_=0.02). Females (M) show increased activity to both CS+ and CS-without discrimination throughout all three days (Figure 4b) (Day 1: F _(1,8)_ = 23.17,n=9, p=0.0013, post hocs: p_CS+_=0.06, p_cs-=_0.012; Day 2: F _(1,8)_ = 30.65,n=9, p=0.0005, post hocs: p_CS+_=0.0007, p_cs-=_0.0028;Day 3: F _(1,8)_ = 23.92,n=9, p=0.0012, post hocs: p_CS+_=0.016,p_cs-=_0.037). (N-Q) Heatmaps illustrating trial by trial fluorescence changes at CS+ and CS-onset in representative animals. (R) Schematic illustrates the use of wavelet signal decomposition to quantify coherence between two fibre photometry time series recorded in mPFC-NAc and vHip-NAc of an individual animal (upper panel). First, the continuous wavelet transform identifies peaks in each time series (middle panel). The product of the two continuous wavelet transforms then generates the cross wavelet transform (lower panel) quantifying coherence between the two signals, with warmer colors indicating higher absolute values. (S) Probing coherence across a range of offsets reveals maximum coherent peaks at 0 sec offset indicating synchronous signals. (T) In males, this synchrony is modulated by aversive cues (F _(2, 12)_ = 9.35, n=7, p=0.004), with a decrease in number of coherent peaks during the CS+ (p=0.02) which returns to higher baseline synchrony during the post-cue period (p=0.01). (U) In females, there is a trending decrease in coherent peaks during the CS+ (F _(2, 13)_ = 6.309, n=9, p=0.012, post-hoc: p=0.08) which is sustained into the post-cue period (p=0.02). In females, but not in males, the CS-exerts an opposing modulation, with an increase in coherent peaks from pre-cue to cue (p=0.02) that returns to baseline in the post-cue period (p=0.004). Graphs show mean +/-SEM.

Time-locked neural activity in both mPFC-NAc and vHip-NAc encoded aversive events and cues predicting these events but discriminating non-threat cues was sex- and pathway-specific. We reasoned that if indeed these pathways carry distinct information between males and females, cue identity should be preferentially recoverable from neural activity in one pathway for each sex. A k-nearest neighbors classifier using late-training cue onset activity achieved reliable classification in females with mPFC-NAc but chance levels with vHip-NAc (Figure S2b,d). Conversely in males, vHip-NAc classification was reliable and mPFC-NAc at chance (Figure S2a,c). Using the full 30sec cue-elicited neural activity increased classifier accuracy across all predictors, however the sex-bias remained with mPFC-NAc outperforming vHip-NAc in females and vHip-NAc outperforming mPFC-NAc in males (Figure S2e-h) confirming sex-specificity in neural discrimination of cues.

mPFC and vHip inputs to NAc are highly convergent ^14^. While examining each pathway in isolation suggested one pathway predominates in each sex, we reasoned that coordinated activity across pathways may also carry information and examined synchrony using wavelet signal decomposition to identify coherent peaks in paired mPFC-NAc and vHip-NAc recordings (Figure 2r). Exploring temporal lag/lead identified maximal coherence at zero offset (i.e. synchrony; Figure 2s) with modulation by cue-type and period. In males, synchrony dropped during CS+ before returning to baseline in the post-cue period (Figure 2t), while in females reduced synchrony during CS+ persisted. Strikingly, in females, but not males, CS-exerted opposing modulation, increasing cue-induced synchrony (Figure 2u). Reduced synchrony may signal a pending aversive event and increased synchrony, resolution of threat or safety. Persistent CS+ induced reduction of synchrony in females may indicate sustained threat perception, defining a background threat-level against which increased CS-induced synchrony can signal safety.

The ultimate output of neural activity is behavior, so we then asked if neural encoding of cue identity drives freezing, a species-specific fear/threat response. Linear mixed effects regression revealed that cue-onset neural activity explains surprisingly little variation in freezing (Table S1, Figure S3). Models integrating both mPFC-NAc and vHip-NAc did not significantly improve prediction (Table S1). Therefore mPFC-NAc and vHip-NAc encode shock-predicting cues, but do not predict freezing, suggesting they do not mediate this behavior.

Analyzing neural activity-behavior relationships indicated that information carried in mPFC-NAc and vHip-NAc does not drive freezing, raising the question of the behavioral significance of cue information in these pathways. Given the role of NAc in integrating cortico-limbic information to regulate motivated behavior ^4,15-17^, we hypothesized that mPFC-NAc and vHip-NAc representations of threat and non-threat cues might modulate reward behavior. To test this, we developed a conditioned suppression paradigm wherein mice learn to suppress rewarded lever pressing during a shock-predicting CS+ (but not CS-) and then chemogenetically inhibited mPFC-NAc or vHip-NAc using an intersectional cre-dependent viral strategy injecting retrograding AAV-cre into NAc and AAV-DIO-hM4Di into mPFC or vHip (Figure 3a-c). Upon stable lever pressing, mice progressed to discriminative fear conditioning with continued reward access and then C21 injected prior to a test with active levers and non-reinforced cues for pathway-specific inhibition (Figure 3c). Inhibiting mPFC-NAc had no effect in males; both hM4Di-DREADD and mCherry-controls suppressed lever pressing to CS+ and not CS-(Figure 3e). In females, mCherry-controls suppressed more to CS+ than CS-, while mPFC-NAc hM4Di-DREADD mice did not discriminate (Figure 3g). With vHip-NAc manipulations, hM4Di-DREADD males showed less suppression to CS+ than mCherry-controls and neither group showed suppression to CS-(Figure 3i). In females, both hM4Di-DREADD mice and mCherry-controls similarly suppressed lever pressing during CS+ and not CS-(Figure 3k). As expected from the minimal prediction of freezing from neural data (Figure S3), freezing was not altered by inhibiting either pathway, with all animals discriminating cues (Figure 3 f,h,j,l). This demonstrates that pathway-specific inhibition leaves cue learning intact but specifically impairs how threat and non-threat cues modulate reward seeking.

**Figure 3.**
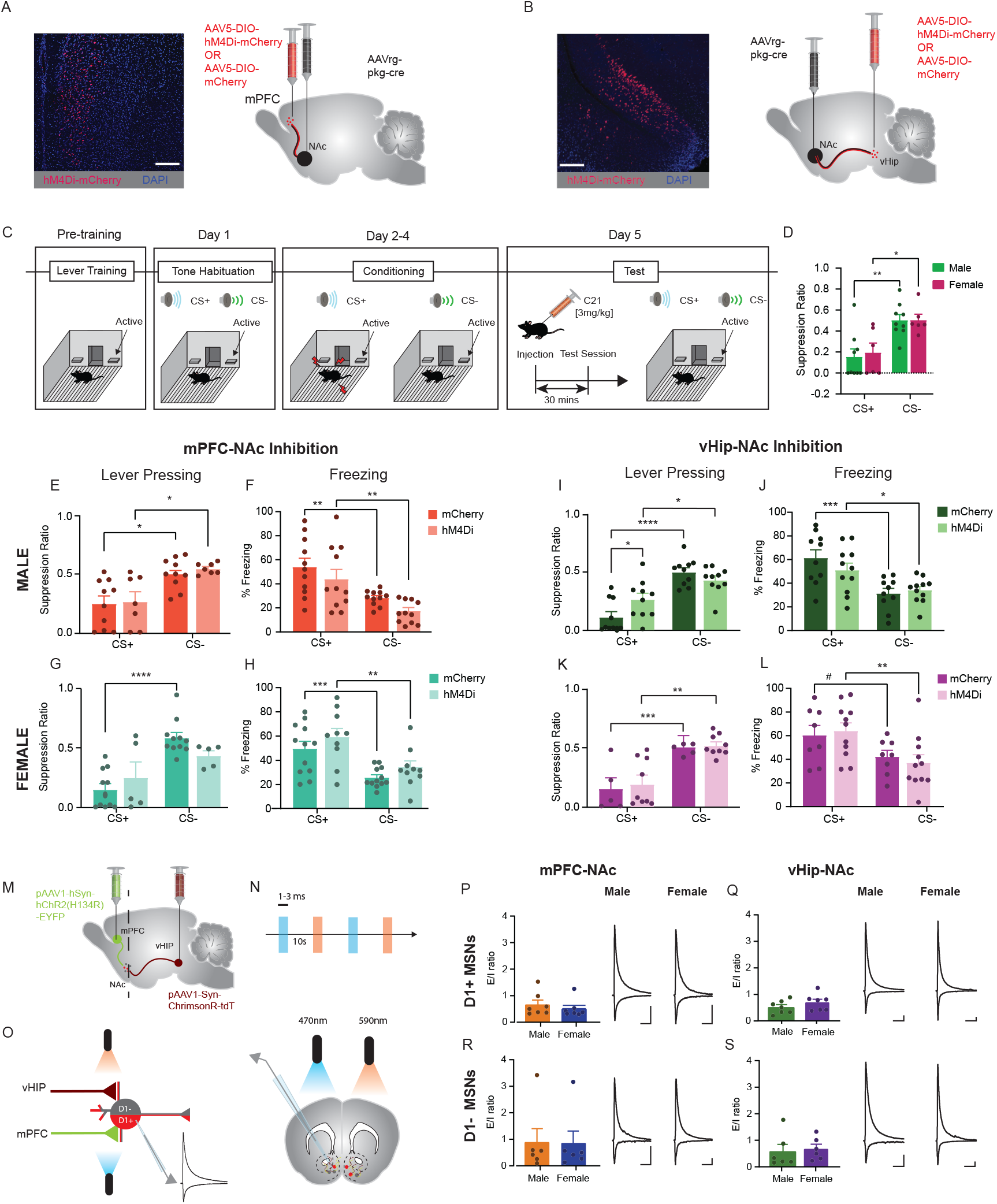
Sexually dimorphic control of reward seeking behavior by accumbal afferents without differences in circuit wiring. Schematics and representative images illustrate an intersectional cre-dependent viral strategy targeted an inhibitory DREADD to either the (A) mPFC-NAc or (B) vHIP-NAc pathway (scale bars represent 200um). (C) Mice were trained in a conditioned suppression paradigm to press a lever for chocolate milk reward prior to a 3-day Pavlovian discriminative fear conditioning paradigm. On a test day, C21 was injected to induce pathway-specific DREADD inhibition and mice were exposed to CS+ and CS-tones without further footshock while lever pressing for reward. (D) Behavioral piloting showed no sex differences in conditioned suppression; both male and female mice showed greater suppression of lever pressing during CS+ versus CS-, indicated by smaller suppression ratios (F _(1,13)_ = 28.77, n=9,6, p=0.0001, *post hocs*: p_M_=0.0012, p_F_=0.012). (E) mPFC-NAc hM4Di-DREADD males showed no differences in suppression ratio (F _(1,15)_ = 18.44,n=10,7, p=0.0006, *post hocs*: p_mCh_=0.012, p_hM4Di_=0.021) compared to mCherry-control males during either cue, and both groups discriminated CS+ and CS-. (F) mPFC-NAc inhibition in males did not impact freezing (F _(1,20)_ = 26.13, n=11,11, p<0.0001, *post hocs*: p_mCh_=0.004, p_hM4Di_=0.004). (G) While female mCherry-control mice suppressed lever pressing more during CS+ than CS-, mPFC-NAc hM4Di-DREADD females did not (F _(1,14)_ = 24.20, n=11,5, p=0.0002, *post hocs*: p_mCh_<0.0001, p_hM4Di_=0.18). (H) mPFC-NAc inhibition also did not modulate freezing in females with both groups freezing more to CS+ than CS-(F _(1,20)_ = 35.12, n=12,10, p<0.0001, *post hocs*: p_mCh_=0.0007, p_hM4Di_=0.0012). (I) vHIP-NAc hM4Di-DREADD males and mCherry-controls discriminated CS+ and CS-, exhibiting smaller suppression ratios (increased suppression of lever pressing) to CS+ than CS-(F _(1,18)_ = 42.2,n=10,10, p<0.0001, *post hocs*: p_mCh_<0.0001, p_hM4Di_=0.021). However, vHIP-NAc hM4Di-DREADD males showed less CS+ mediated suppression than mCherry-controls (F _(1,18)_ = 6.103, n=10,10, p=0.024, *post hocs*: p=0.039. (J) vHip-NAc inhibition did not alter cue discrimination assessed by freezing (F _(11,,18)_ = 26.12, n=10,11, p=0.0005, *post hocs*: p_mCh_<0.0001, p_hM4Di_=0.03). (K) Female vHip-NAc hM4Di-DREADD and mCherry-controls both discriminated CS+ and CS-with no group differences in suppression ratio (F _(1,12)_ = 54.36, n=5,9, p<0.0001, *post hocs*: p_mCh_=0.0077, p_hM4Di_=0.0009). (L)There were also no group differences in freezing during the CS+ cue (F _(1,17)_ = 22.96, n=8,11, p=0.0004, *post hocs*: p_mCh_=0.067, p_hM4Di_=0.0017). (M) To probe circuit connectivity, animals were bilaterally injected with AAV-ChR2 and AAV-ChrimsonR in mPFC and vHIP, respectively. (N,O) 470nm and 590nm light (1-3ms) was alternated to stimulate mPFC and vHIP terminals, respectively, during patch clamp recordings of optical EPSCs and IPSCs in D1+ and D1-cells in the NAc medial shell. Bar graphs and representative traces show no differences were observed in oEPSC/oIPSC ratio of D1-MSNs evoked by either (P) mPFC (t_12_=0.75, n=7,7, p=0.46) or (Q) vHIP (t_12_=1.88, n=7,7, p=0.26) or D2-MSNs evoked by either (R) mPFC (t_10_=0.07, n=6,6, p=0.94) or (S) vHIP (t_10_=0.27, n=6,6, p=0.79) (scale bars represent 100ms on the x-axis and 100pA on the y-axis). Graphs show mean +/-SEM.

vHip-NAc inhibition attenuated conditioned suppression in males but not females whereas mPFC-NAc inhibition impaired conditioned suppression and cue discrimination in females but not males. mPFC and vHip inputs converge in NAc medial shell to regulate medium spiny neuron (MSN) firing ^18,19^. Contrasting effects of pathway-specific inhibition between males and females could arise from sex differences in behavioral strategy mediating differential recruitment of similarly wired circuits, or alternatively, sex differences in circuit connectivity ^20,21^. We therefore probed possible sex differences in synaptic drive onto the two populations of NAc MSNs, Drd1-enriched (D1-MSNs) and Drd2-enriched (D2-MSNs; largely Drd1-negative). We injected D1-tdTomato mice with AAV-ChR2 in mPFC and AAV-ChrimsonR in vHip to independently stimulate each pathway (Figure 3m), prepared acute NAc slices and recorded D1+ and D1-(putative D2+) MSNs (Figure 3n,o). Optogenetically-evoked EPSCs (oEPSCs) and IPSCs (oIPSCs) were successfully evoked from both inputs in all recorded post-synaptic cells (alternating blue/orange light; Figure 3n,o) confirming widespread single-cell input convergence. There were no sex differences in oEPSC/oIPSC ratio for mPFC or vHip in D1+ or D1-MSNs (Figure 3p-s) confirming similar drive on both cell-types from both pathways in males and females. This strongly suggests that sex differences in pathway-specific behavioral control do not result from underlying differential circuit connectivity but rather from differential recruitment of similarly wired circuits in males and females as expected from the sex-specific neural encoding revealed by fibre photometry (Figure 2).

Integration of reward and aversion is central to adaptive behavior and dysregulated in various psychopathologies ^22^. Surprisingly, despite similar behavior and circuit connectivity, we identify a sexual dimorphism in the neural encoding of threat discrimination that may indicate sex-specific strategies in the face of threat. We find that vHip-NAc, in males, and mPFC-NAc, in females, encode aversive cues and discriminate these from neutral cues to modulate reward behavior under threat. Modest differences in freezing as well as alterations in inter-pathway synchrony suggest that females, but not males, process a non-reinforced cue as a safety signal, which may contribute to differential pathway recruitment. Sex-specificity in threat processing may underlie sex differences in vulnerability to stress-related disorders given that chronic stress differential impacts transcription and morphology across different brain regions in males and females^23,24^. Here we identify novel mechanisms of threat discrimination and behavioral integration in male and female mice, demonstrating that similar behavior need not equal similar mechanism.

## Methods

### Animals

Mice were maintained on a 12-h light-dark cycle (lights on at 7:00AM) at 22-25°C group-housed with 2-3 same-sex cage-mates with *ad libitum* access to food and water. All experimental manipulations occurred during the light cycle, in accordance with guidelines of McGill University’s Comparative Medicine and Animal Resources Center and approved by the McGill Animal Care Committee.

Neural Recordings: D1-cre mice were used with an initial plan to simultaneously interrogate activity in the NAc, but due to technical difficulties, this data was discarded. D1-cre heterozygote mice bred on C57BL/6J background were obtained from Jackson laboratories and bred with D1-cre wild type mice. Pups were weaned at post-natal day 21 and housed with same-sex litter mates and heterozygote mice were separated 4 weeks post-weaning and housed with 2-3 same sex littermates.

Chemogenetic/optogenetic manipulations: 7-week-old male and female C57BL/6J mice were obtained from Jackson Laboratories and habituated to the colony room one week prior to start of manipulations. Mice were food restricted to 85% of their free-feeding body weight during experimentation.

Ex vivo electrophysiology: Male and female D1-tdTomato BAC transgenic mice (B6.Cg-Tg(Drd1a-tdTomato)6Calak/J) initially obtained from Jackson Laboratories were bred at the Comparative Medicine and Animal Resources Centre (CMARC) at McGill University.

### Surgeries

Stereotaxic surgery was performed under ketamine (100 mg/kg)/xylazine (10 mg/kg) anesthesia. To achieve projection-specific GCaMP7f expression in glutamatergic NAc-projecting cells, 0.5μl pGP-AAVrg-syn-jGCaMP7f-WPRE virus ^25^ (1.85× 10^13^GC/ml; Addgene) was infused into the NAc (A/P: +1.3, M/L: +/-0.60, D/V: −4.9) at a rate of 0.1μl per min, before raising the needle to D/V: −4.7 and infusing a further 0.5µl virus, allowed to diffuse for 10 min before withdrawing the needle. Chronically implantable optic fibers (Neurophotometrics) with 200μm core and 0.37 NA threaded through ceramic ferrules were implanted above the ventral subiculum of the vHip (A/P: −3.40, M/L: +/-3.00, D/V: −4.75) and infralimbic mPFC (A/P: −0.3, M/L: +/-1.90, D/V: −2.80). pGP-AAV-syn-jGCaMP7f-WPRE was a gift from Douglas Kim & GENIE Project (Addgene plasmid # 104488; http://n2t.net/addgene:104488; RRID:Addgene_104488). Recordings began minimum 4 weeks after surgery to allow sufficient time for stable and robust retrograde virus expression.

To achieve projection specific expression of inhibitory designer receptors exclusively activated by designer drugs (DREADD) for chemogenetic manipulation, 0.5 μl of AAV-rg-pkg-cre (1.17 × 10^13 GC/ml; Addgene); diluted with sterile PBS at a ratio of 1:4 was bilaterally injected into the NAc (AP:+1.30, ML: +/-1.81, D/V: −4.43, 15° angle) and 0.5 μl of AAV5-hSyn-DIO-hM4D(Gi)-mCherry (2.4 × 10^13 GC/ml; Addgene) ^26^ or AAV5-hSyn-DIO-mCherry (2.3 × 10^13 GC/ml; Addgene) was bilaterally injected into either the mPFC (AP:1.7, ML: +/-0.75, D/V: −2.5, 12° angle) or vHip (AP:-3.40, ML: +/-3.95, D/V: −4.17, 12° angle). AAV-pgk-Cre was a gift from Patrick Aebischer (Addgene plasmid # 24593; http://n2t.net/addgene:24593 ; RRID:Addgene_24593). pAAV-hSyn-DIO-hM4D(Gi)-mCherry was a gift from Bryan Roth (Addgene plasmid # 44362; http://n2t.net/addgene:44362 ; RRID:Addgene_44362). pAAV-hSyn-DIO-mCherry was a gift from Bryan Roth (Addgene plasmid # 50459 ; http://n2t.net/addgene:50459 ; RRID:Addgene_50459). Coordinates were adjusted to facilitate simultaneous bilateral injections targeting the same subregions as for fibre photometry. Manipulations began a minimum of 4 weeks after surgery to allow sufficient time for stable and robust retrograde virus expression.

For ex vivo electrophysiology experiments, stereotaxic injections were conducted on P56 – P77 mice. AAV vectors expressing ChrimsonR (pAAV1-Syn-ChrimsonR-tdT; 2 × 10^13 GC/ml; Addgene) ^27^ or channelrhodopsin-2 (hChR2) (pAAV1-hSyn-hChR2(H134R)-EYFP; 2.1 × 10^13 GC/ml; Addgene) were used for independent optical stimulation of glutamatergic afferents in the NAc shell from the ventral hippocampus (vHip) and the infralimbic mPFC, respectively. pAAV-Syn-ChrimsonR-tdT was a gift from Edward Boyden (Addgene plasmid # 59171 ; http://n2t.net/addgene:59171 ; RRID:Addgene_59171). pAAV-hSyn-hChR2(H134R)-EYFP was a gift from Karl Deisseroth (Addgene plasmid # 26973 ; http://n2t.net/addgene:26973 ; RRID:Addgene_26973). Virus was bilaterally infused 0.5 μl pAAV1-hSyn-hChR2(H134R)-EYFP at a rate of 0.1 μl / min into infralimbic mPFC and 0.5 μl pAAV1-Syn-ChrimsonR-tdT into vHip.

### Perfusions and Histology

Mice were anesthetized with ketamine (100 mg/kg)/xylazine (10 mg/kg) and transcardially perfused with 1xPBS and formalin. Brains were post-fixed in formalin for 24hrs before being transferred to PBS. Brains were sectioned on a vibratome at 50µm. Sections were mounted and coverslipped with Vectashield (Vectorlabs). Histology was confirmed on an epifluorescent microscope (Leica) to verify viral expression in targeted regions and accuracy of fiber placement where applicable. Mice were excluded for lack of viral expression, off-target expression or misalignment of virus and fiber. One mouse with misaligned fiber/virus in vHIP and another with bilaterally mistargeted DREADD expression were excluded from data analysis.

### Apparatus

Behavioral experiments were performed in standard MED Associates operant boxes (15.24 × 13.34 × 12.7 cm) enclosed in sound attenuating chambers outfitted with a programmable audio generator and a house light to deliver cues, shock-enabled grid floors and two retractable levers either side of a food port for delivering liquid chocolate milk reward (Nesquick). Food ports were closed off during Pavlovian fear conditioning. Data was collected and protocol was run using MED-PC software. Behavior was recorded using Raspberry Pi cameras for offline analysis.

### Pavlovian Fear Conditioning

Following a 120s habituation period, mice were exposed to 8, 30s presentations each of an auditory cue (2000Hz, 63dB) and a visual cue (house light). One cue served as a conditioned threat cue (CS+) and co-terminated with a 0.5s, 0.5mA shock, while the other as an unconditioned cue (CS-) with no outcome; cue identity was fully counterbalanced. Cues were randomly presented and were followed by a variable intertrial interval (ITI) averaging 90s.

Freezing ratio (FR) was calculated as follows:

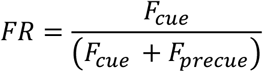

Where F is the percent time freezing during the cue and F is the percent time freezing during an equivalent pre-cue period. A freezing ratio of 0.5 indicates equal freezing during cue and pre-cue. FR greater than 0.5 indicates increased freezing during the cue while an FR less than 0.5 indicates a reduction in freezing.

Discrimination score (DS) was calculated as follows:

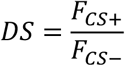

Where F_CS+_ and F_CS-_ are the percent time freezing during the CS+ and CS-cues respectively.

### Frame Independent Projected Fiber Photometry

To measure calcium-associated changes in fluorescence in real time, recordings were made from vHip-NAc and mPFC NAc-projecting cells during discriminative Pavlovian fear conditioning. Samples were collected at a frequency of 20 Hz using Neurophotometrics hardware through Bonsai and FlyCap software. Recordings were coupled to the start of behavioral analysis by interfacing Bonsai with MED-PC using a custom DAQ box (Neurophotometrics). Data were extracted and analyzed using custom-written scripts in Matlab R2019a (The MathWorks) ^28^. To normalize the data and control for potential artifacts, the control channel (415nm) was fitted to the raw (470nm). The fitted control was then subtracted from the raw trace. The resultant trace was divided by the fitted control giving the ΔF/F and converted to a z-score. The subsequent trace was detrended to remove bleaching artifacts ^29^.

### Cross Wavelet Transform

To quantify the degree of coherence between vHIP-NAc and PFC-NAc pathways, we employed a cross wavelet transform (XWT) using the Python package PyCWT ^30^. Briefly, a continuous wavelet transform (CWT), using the Mexican hat wavelet, was applied to the neural traces from both regions to detect peaks in activity in three separate windows: a pre-cue period ranging from 29 secs before cue onset to 2 secs before cue onset, a cue period ranging from cue onset to 3 secs before cue offset and a post-cue period ranging from 2 secs following cue offset to 29 secs following cue offset. The period around cue offset was excluded to remove the very large increases in activity following footshock. After calculating the CWTs the product of the two CWTs was taken to generate the XWT, from which synchronous events were identified as follows. At each time point, the maximum value of the XWT over frequency range 0.5Hz to 1.0 Hz was found. For the resulting waveform, peaks with a significance value exceeding 0.9 ^30^ that were the product of two positive CWT values were classed as synchronous events. This process was also performed using offset CWTs for time offsets ranging from −1 sec to +1 sec from 0, to identify the offset between signals at which the highest degree of coherence was observed, with zero offset indicating peaks that are synchronous between the two pathways, and non-zero offsets indicating lag/lead. We compared the number of synchronous peaks (within a 0.1sec window) during pre-cue, cue and post-cue periods.

### Conditioned Suppression

Mice were presented with 2 extended levers with a press on the active lever reinforced by a liquid reward (Nesquik chocolate milk, diluted with water at a ratio of 2:1) and presses on the inactive lever followed by no outcome. Mice started on a fixed ratio 1 (FR1) delivery schedule then progressed through random ratio (RR) training with 20% (RR5), 10% (RR10) and then 5% (RR20) of active lever presses rewarded. Following RR20, mice were trained on a modified Pavlovian fear conditioning with continued reinforced RR20 reinforced lever pressing. Prior to conditioning, mice went through one session of tone habituation in which they were exposed to 6, 30sec presentations each of two tones (A; 10000Hz, 72 dB; B: 2000Hz, 63dB) with no outcomes. Conditioning consisted of 6 presentations of these same cues counterbalanced as either CS+ or CS-cues; CS+ cue co-terminated in a 0.5mA, 0.5sec shock, CS-no outcome. Following three days of training, conditioned suppression was tested under fear extinction conditions with levers continuing to be reinforced. Mice performing less than 100 lever presses during test were excluded from analysis. Number of excluded mice di not significantly differ between experimental groups. Custom scripts in combination with EZ track software were used to quantify freezing behavior.

Suppression ratio (SR) was calculated as follows:

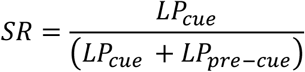

Where LP_cue_ is the number of lever presses during the cue and LP_pre-cue_ is the number of lever presses during an equivalent pre-cue period. A suppression ratio of 0 indicates complete suppression and a ratio of 0.5 indicates no suppression.

### Chemogenetic Manipulations

Compound 21(C21) (Hello Bio) was dissolved in 0.9% NaCl and stored in 2mL vials at −20°C. A new vial was thawed before use every day. 30 mins before the start of the test session, mice received an intraperitoneal injection of Compound 21 (C21) (3mg/kg) to activate inhibitory DREADDs.

### Ex vivo patch clamp electrophysiology

Brain slice preparation: Mice were deeply anesthetized with isofluorane. Transcardial perfusion was performed with 20-25 ml of ice-chilled carbogenated NMDG artificial cerebrospinal fluid (ACSF) (containing in mM: 92 NMDG, 2.5 KCl, 1.25 NaH_2_PO_4_, 30 NaHCO_3_, 20 HEPES, 25 Glucose, 2 Thiourea, 5 Na-ascorbate, 3 Na-pyruvate, 0.5 CaCl_2_·4H_2_O and 10 MgSO_4_·7H_2_O; pH: 7.3–7.4; osmolality: 300-310 mOsmol/kg). Brain slices (200 *μ*m) were prepared in ice-chilled carbogenated NMDG aCSF by a vibratome (Lecia VT 1200S). After slice preparation, all brain slices were recovery in 32–34 °C carbogenated NMDG ACSF for 10 min. Then, brain slices were transferred into room-temperature carbogenated HEPES holding aCSF (containing in mM: 92 NaCl, 2.5 KCl, 1.25 NaH_2_PO_4_, 30 NaHCO_3_, 20 HEPES, 25 Glucose, 2 Thiourea, 5 Na-ascorbate, 3 Na-pyruvate, 2 CaCl_2_·4H_2_O and 2 MgSO_4_·7H_2_O; pH: 7.3–7.4; osmolality: 300-310 mOsmol/kg). Brain slices were kept in HEPES holding ACSF before recording.

Recordings: Whole-cell recordings were performed in room-temperature carbogenated ACSF (containing in mM: 128 NaCl, 3 KCl, 1.25 NaH_2_PO_4_, 2 MgCl_2_, 2 CaCl_2_, 24 NaHCO_3_ and 10 Glucose, pH 7.2; osmolality: 300-310 mOsmol/kg). Patch pipettes were filled by cesium-methanesulfonate based internal solution (containing in mM: 130 Cs-methanesulfonate, 10 HEPES, 0.5 EGTA, 8 NaCl, 5 TEA-Cl, 4 Mg-ATP, 0.4 Na-GTP, 10 Na-phosphocreatine and 1 QX-314; pH 7.2; osmolality: 290-300 mOsmol/kg) for voltage-clamp recordings. D1+ medium spiny neurons (MSNs) and D1-MSNs were patched to record excitatory / inhibitory (E/I) ratio. MSNs were held at −70 mV to measure α-amino-3-hydroxy-5-methyl-4-isoxazolepropionic acid receptor (AMPAR)-mediated currents in the present of *N*-methyl-D-aspartate receptor (NMDAR) antagonist, 3-(2-carboxypiperazin-4-yl) propyl-1phosphonic acid (CPP, 10 mM; HelloBio). Inhibitory postsynaptic currents (IPSCs) were recorded at AMPAR reversal potential (CPP, 10 mM) at +20 mV.

In every patched MSN (D1+ or D1-), E/I ratio was probed by the activation of afferents from both the IL-PFC and the vHip. Optically evoked EPSCs and IPSCs were obtained every 10secs, alternating pathways. Two different wavelengths of lights from a LED system (DC4100, Thorlabs) were used to activate afferent fibers from the IL-PFC (wavelength: 470 nm) and the vHip (wavelength: 590 nm) in alternation. A pilot study established optimal parameters for independent pathway stimulation. Light pulse width (1-3 ms) and light intensity (150 pA) for both wavelengths of light were held constant for each neuron. This yielded stable EPSCs (50 – 2000 pA) at −70 mV. All signals were amplified and digitized by Multiclamp 700B (Molecular Device) and Digidata 1550B (Molecular Device), respectively. Series and access resistance were monitored during the experiments and signals were bessel filtered at 2 kHz.

### Statistics

Inferential statistical analyses were performed using GraphPad Prism 7. Grubb’s test was used to identify and exclude statistical outliers. One sample two-tailed t-tests were used to assess deviation of sample means from a baseline. Paired, two-tailed, two sample t-tests were used to compare means of two within-group repeated-measures. Unpaired, two-tailed, two-sample t-tests were used to compare two means of between-group measures. Two-way repeated measures ANOVAs with Sidak’s multiple comparisons post-hoc tests were used to compare between and within group measures where two groups were tested on two or more measures.

Linear mixed effects regressions were run in R with neural activity at cue onset as a fixed effect, an arbitrary categorical variable assigned to each animal to account for inter-individual variability as a random effect and freezing as the dependent variable. An ANOVA was run to compare models using either vHip-NAc, mPFC-NAc or both vHip-NAc and mPFC-NAc activity at cue onset as a fixed variable and inter-animal variability and a random variable to a null model which includes only inter-animal variability as a predicting variable. Chi squared and p-values for the ANOVA are shown in table S1. Marginal R^2^ (R^2^m) represent the variability explained by fixed effects while conditional R^2^ (R^2^c) represents the variance explained by the entire model.

Generalized additive models (GAMs) were generated using the “bam” function in R with an arbitrary categorical variable assigned to each animal to account for inter-individual variability as a random effect and cue type as a fixed effect. We generated a set of smooth functions to capture contributions of cue type to variation in neural activity and took the difference of CS+ and CS-functions to find periods of maximal difference in neural encoding.

K Nearest Neighbor classifier was run using KNeighborsTimeSeriesClassifier from tslearn library. Time series containing delta FF from either mPFC-NAc recordings or vHip-NAc recordings from either the first second of the cue or the entire cue period were used as input for the classifier and labeled based on cue type. Number of nearest neighbors (K) was made a hyperparameter.

## Acknowledgements

This work was supported by funding from the Ludmer Centre for Neuroinformatics & Mental Health and a CIHR Project grant (201709PJT-391173-BSA-CFAA-178116 to RCB, and a CIHR graduate scholarship (201810GSD-4221 05-DRA-CFAA-297096) to JM. Images were collected in the McGill University Advanced BioImaging Facility (ABIF), RRID:SCR_017697.

## Ethics Statement

All authors declare no conflicts of interest.

