## Supplemental Figures for "Sexually dimorphic neural encoding of threat discrimination in nucleus accumbens afferents drives suppression of reward behavior"

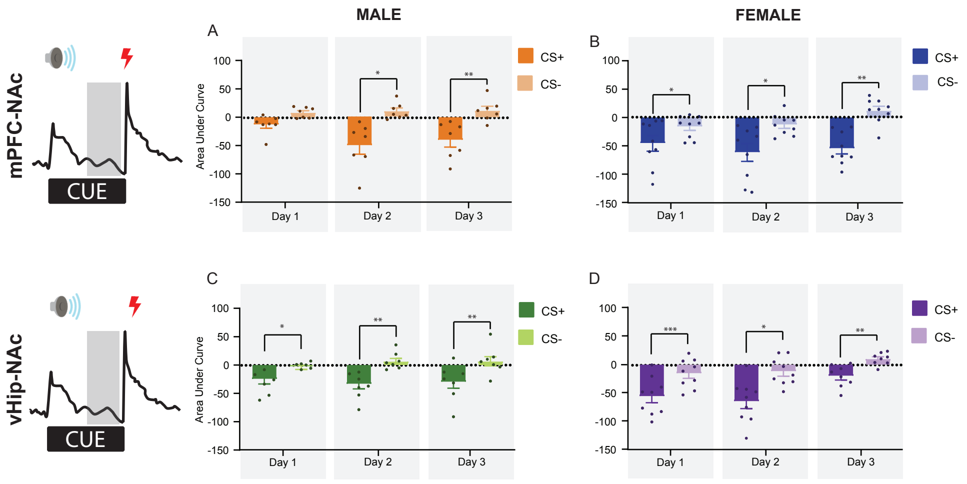


Supplemental Figure 1. CS+ mediates pre-outcome suppression in neural activity. PFC-NAc activity is suppressed during the final 8 secs of the CS+ compared to CS-, with a trend in (A) males(n=8, p=0.068) and a significant decrease in (B) females in early training (n=9, p=0.02) which rises to significance in mid (M: n=8, p=0.027, F: n=9, p=0.04) and late training (M: n=7, p=0.0041, F: n=9, p=0.0016) in both sexes. vHIP-NAc activity is suppressed in males (Day 1: n=6,p=0.01; Day 2: n=7, p=0.0035; Day 3: n=7, p=0.02) and females (Day 1: n=9 p=0.0002; Day 2: n=9 p=0.019; Day 3: n=9, p=0.034) throughout all days of training. Data represented as mean +/- SEM.

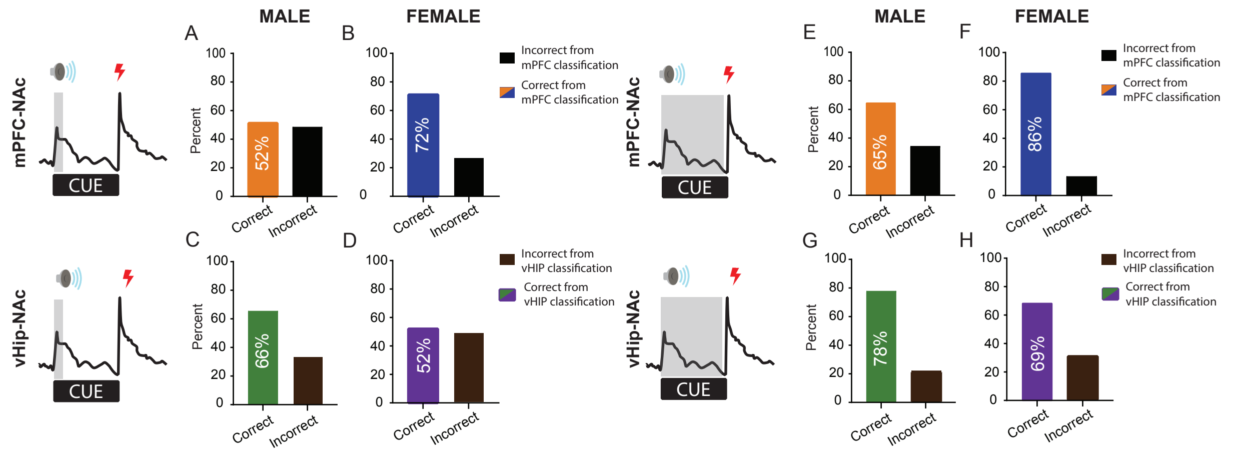


Supplemental Figure 2. Accumbal afferents encode cue type in a sex-specific manner. Using a K-Nearest Neighbor's (KNN) classifier, cue identity was predicted from neural activity during the first 1 sec (A-D) or entirety (E-H) of the cue. mPFC-NAc activity classified cue identity at chance (52%) in males (A) but with 72% accuracy in females (B). Conversely, vHIP-NAc classification was reliable in (C) males (68%) but at chance levels in (D) females (52%). Using the full 30 sec of cue-elicited neural activity increased classifier accuracy across all predictors, however a sex-bias remained with (F) mPFC-NAc activity retaining reduced accuracy in males (65%) compared to female (86%) and vHIP-NAc (69%) maintaining increased accuracy in (C) males (78%) compared to in (D) females (69%).

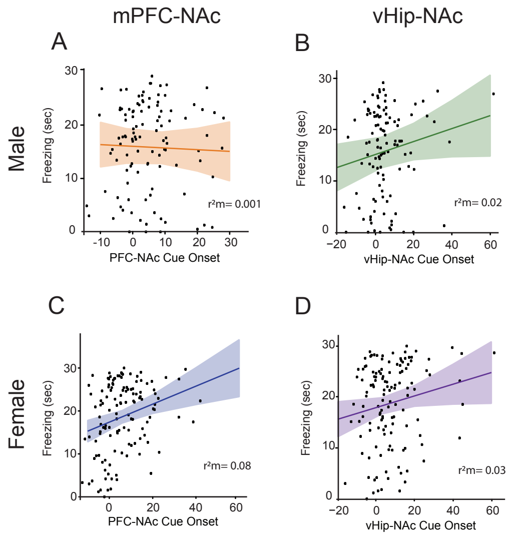


Supplemental Figure 3. Accumbal afferent activity explains little variability in freezing behavior. A linear mixed effects regression assessed the relationship between freezing and neural activity while controlling for inter-individual variability. In males, (A) mPFC-NAc activity accounts for less than 1% of variance in freezing, while (B) vHIP-NAc activity at cue onset accounts for 2.5%. In females, (C) mPFC-NAc activity accounts for 8% of variance while (D) vHIP-NAc activity accounts for 3%. Plots show regression line +/- 95% confidence interval.

| | Fixed variable | AIC | BIC | Log(Lik) | deviance | $R^2$m | $R^2$c | ChiSq | P |
|---|---|---|---|---|---|---|---|---|---|
| **Female** | None | 903.26 | 911.8 | -448.6 | 897.26 | NA | 0 | | |
| | vHip Cue Onset | 901.36 | 912.8 | -446.7 | 893.36 | 0.03 | 0.04 | 9.079 | 0.05 * |
| | mPFC Cue Onset | 894.15 | 905.55 | -443.1 | 8886.2 | 0.08 | 0.1 | 11.118 | 0.0009* |
| | mPFC+vHip Cue Onset | 895.82 | 910.08 | -442.91 | 885.82 | 0.08 | 0.1 | 11.441 | 0.003* |
| **Male** | None | 794.62 | 802.77 | -394.31 | 788.62 | NA | 0.15 | | |
| | vHip Cue Onset | 793.23 | 804.10 | -392.61 | 785.23 | 0.03 | 0.18 | 0.1598 | 0.69 |
| | mPFC Cue Onset | 796.46 | 807.33 | -394.23 | 788.46 | 0.001 | 0.15 | 3.3891 | 0.06 |
| | mPFC+vHip Cue Onset | 792.52 | 806.12 | -391.26 | 782.52 | 0.06 | 0.20 | 6.0909 | 0.05 * |

Supplementary Table 1. Relationship between neural activity at cue onset and freezing behavior. ChiSquare and p values for comparisons of model with no fixed variable (random variable only) and a model with the indicated variable as a fixed variable as well as a random variable (see methods for detail). In females, vHIP-NAc activity at cue onset and mPFC-NAc activity at cue onset significantly improve a null model and explain 3% and 8% of variance, respectively. In males, neither vHIP-NAc or mPFC-NAc activity at cue onset improve a null model and explain 0.1% and 3% of variance, respectively.
